# Engineered microvasculature using maskless photolithography and on-chip hydrogel patterning: a facile approach

**DOI:** 10.1101/2024.07.22.604661

**Authors:** Dhanesh G. Kasi, Mees N. S. de Graaf, Dennis M. Nahon, Francijna E. van den Hil, Arn M. J. M. van den Maagdenberg, Christine L. Mummery, Valeria V. Orlova

## Abstract

*In vitro* models of human microvasculature are increasingly used to understand blood vessel diseases and to support drug development. Most engineered models, however, are slow and labor-intensive to produce. Here, we used a single commercial digital micromirror device (DMD)-based setup for maskless photolithography to both fabricate microfluidic chips and pattern the inside of these chips with gelatin methacrylate (GelMA) hydrogels. These hydrogel scaffolds had tunable stiffness, could be generated rapidly and were suitable for forming perfusable microvasculature from human induced pluripotent stem cell-derived endothelial cells (hiPSC-ECs). When cultured in narrow channels, the hiPSC-ECs adopted a tubular morphology that was similar to capillaries *in vivo*, but they followed the square channel geometry in wider channels. Compartmentalization of the chips allowed co-culture of hiPSC-ECs with hiPSC-derived astrocytes, thereby increasing model complexity. Furthermore, valve-like structures could be patterned inside the channels, mimicking functional vascular valves, holding promise for thrombosis and lymphatic vasculature research.

## Introduction

The microvasculature is integral to the vascular system. It plays a crucial role in maintaining tissue homeostasis by facilitating nutrient and oxygen exchange and removing waste. It is also a primary route for drug delivery and is thus an important target for therapeutic interventions^1–3^. *In vitro* models of human microvasculature are increasingly popular and regarded as valuable tools for studying vessel diseases, drug development, and tissue vascularization. The recent use of human induced pluripotent stem cells (hiPSCs) as a source of vascular cells in particular, allows the generation of disease- and patient-specific vasculature, enhancing the relevance of these models for human health^4–8^.

Conventional two-dimensional (2D) vascular models have been useful to study endothelial cells (ECs) but lack complexity and physiological relevance. hiPSC-based three-dimensional (3D) models provide opportunities to solve these issues while also offering increased throughput and relevance compared to animal models^5,9,10^. Considerable effort has thus been directed towards engineering advanced models that replicate essential features of human microvasculature^4,7,11^.

Notably, microfluidics has been a significant driver in advancing the development of microphysiological systems (MPS), given that it enables the formation of perfusable microvasculature *in vitro*^4,10,12^. Self-assembled microvascular networks can be formed in microfluidic chips because of the intrinsic ability of ECs to self-organize. However, self-organization does not allow precise control of the vessel geometry, resulting in variability in vessel dimensions^12–16^. To overcome this limitation, 3D bioprinting has emerged as a promising technique that enables fabrication of well-defined, uniform vascular constructs. However, integrating 3D printing with microfluidics has not been straightforward. The use of “open-top” chips with a relatively large compartment for printer access is usually required, and, depending on the specific model being developed, may also require sealing the chip after printing. Furthermore, the smallest features achievable are typically greater than 100 µm due to limited printing resolution and the process is time-consuming, requiring “layer-by-layer” fabrication^17–22^. Other fabrication techniques, such as two-photon polymerization (2PP), support enhanced high-resolution manufacturing of 3D structures and are based on both semi-synthetic and synthetic hydrogels, thereby facilitating straightforward patterning inside microfluidic chips. Additionally, innovative methods such as direct laser patterning of hydrogels, including photoablation and cavitation molding, facilitate the precise formation of microchannels within native hydrogels such as collagen, eliminating the need for synthetic hydrogels. The high precision of these methods offers a significant advantage by allowing more controlled structure creation but comes at the cost of limited throughput, as high precision necessitates slower processing speed^23–32^. On the other hand, more traditional methods, such as micromolding and template casting, which are also compatible with native hydrogels, have produced valuable vascular constructs. However, these methods are challenging to scale, are labor-intensive and difficult to perform directly on-chip^33–37^.

Lithography-based manufacturing methods such as 2PP and bioprinting based on digital light processing (DLP) require hydrogel precursors with appropriate photochemistry and a photoinitiator to produce photo-crosslinked hydrogels. Beyond its application in bioprinting, DLP is used in maskless photolithography. The technology is based on a digital micromirror device (DMD), which spatially modulates light to project arbitrary digital photomasks, thereby allowing rapid microfabrication^38–43^. Additionally, when integrated with an inverted microscope, DMD-based systems enable patterning of cells, polymer brushes, proteins and hydrogels^44–51^. However, systems like this are often custom-built in-house and not easily accessible to most researchers. The PRIMO system (Alvéole, France), by contrast, is available as a ready-to-use DMD-based system for maskless photolithography which has been used in several different applications previously and can be implemented in facile bioengineering workflows^42,43,52–55^.

Here, we show how this maskless photolithography system can be used for direct hydrogel patterning on-chip to fabricate various perfusable microvascular structures and microscopic valves. The PRIMO system was utilized for both the initial cleanroom-free chip fabrication (as we described earlier^42^), and subsequent on-chip hydrogel patterning. By using gelatin methacrylate (GelMA) and lithium phenyl-2,4,6-trimethylbenzoylphosphinate (LAP) as the photoinitiator, we were able to pattern microvascular scaffolds and structures rapidly, directly on-chip. GelMA is a gelatin-based hydrogel commonly used for tissue engineering and 3D bioprinting applications. The functionalization of methacrylate allows crosslinking by a photoinitiator and an appropriate light source^23,56–59^.

The patterned microvascular scaffolds could be seeded with hiPSC-derived ECs (hiPSC-ECs) and the resulting vasculature ranged in size from arteriole-sized microvessels (100 µm) to tubular capillary-sized structures with diameters as small as 10 µm. Moreover, hydrogel stiffness, swelling and compliance could be easily controlled by changing the laser dose, adding an additional level of tunability. By simply controlling the hydrogel pattern and adding additional outlets to the gel chamber, we introduced compartmentalization and were able to add hiPSC-derived astrocytes embedded in a fibrin gel. This allows multicellular interactions to be studied and enables modeling of more complex structures such as the blood-brain barrier. Finally, we patterned valve-like structures with leaflets inside the hydrogel channels and demonstrated that advanced microvasculature with functional valves could be generated. Valved microvessels can be used to model and study thrombosis, associated processes (e.g. vortical flow and platelet aggregation) and lymphatics *in vitro*. Taken together, the level of control shown here allows rapid engineering of a wide range of different types of perfusable microvasculature and provides opportunities for vascular disease modeling.

## Results and Discussion

### Hydrogel patterning inside microfluidic chips

Using the PRIMO system, microfluidic chips with a height of 50 µm were fabricated for subsequent hydrogel patterning on the inner surface (Fig 1A and B). Replicating the glass SU-8 mold using an epoxy-resin enhanced durability and increased the number of molds that could be produced (Figure 1B). The PRIMO system allows projection of any chosen pattern as a digital photomask using a DMD, UV light source (375 nm), and microscope optics. When required, several Fields-of-View (FoVs) can be stitched together to project large continuous patterns or arrays onto the substrate. Microfabrication and hydrogel patterning can thus be carried out using a single setup^42,43,53^.

**Figure 1.**
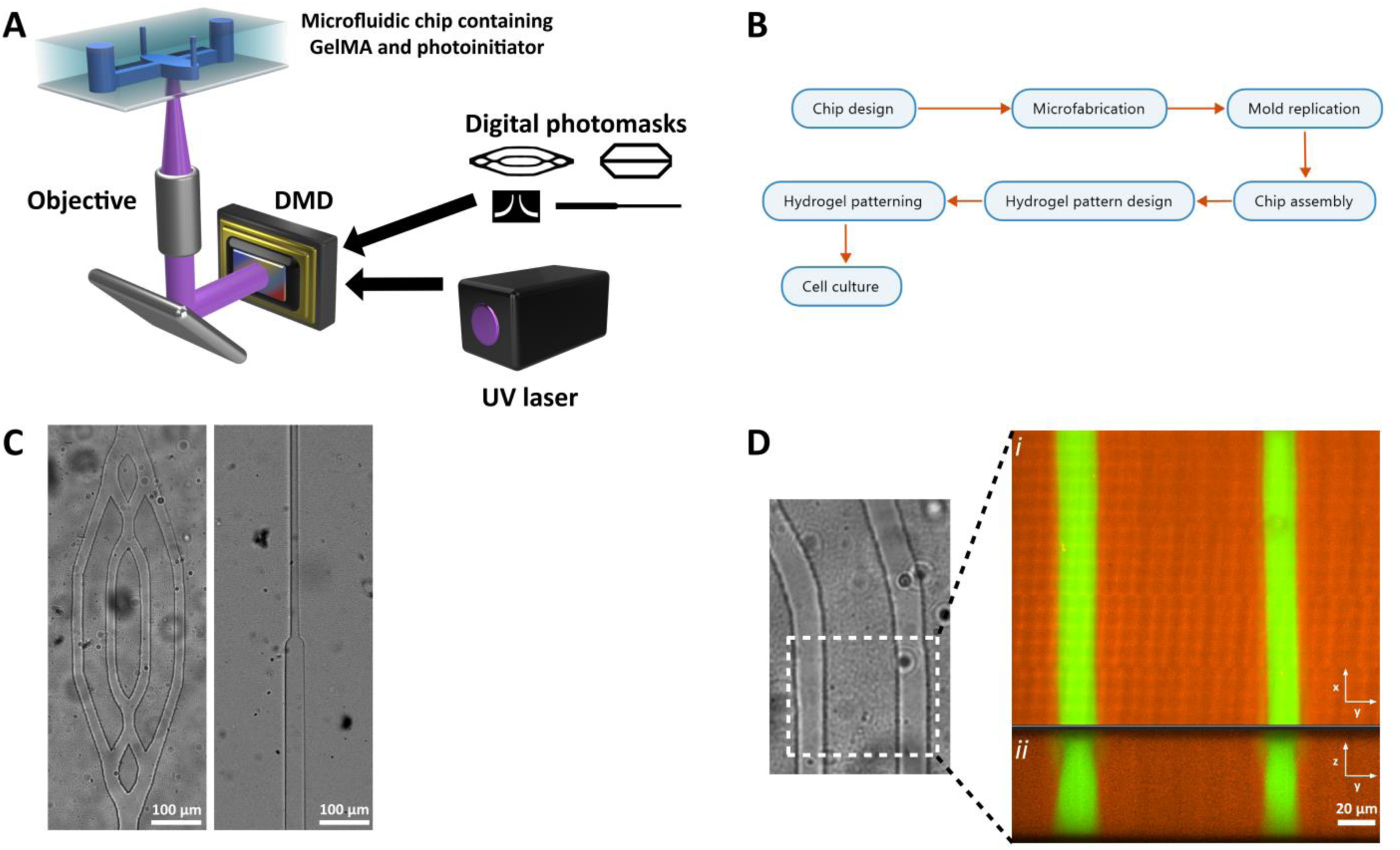
Hydrogel patterning inside microfabricated chips. **(A)** Hydrogel patterning was performed using the PRIMO system. A DMD spatially modulates UV light and projects arbitrary digital photomasks onto the microfluidic chip using conventional microscope optics, thereby crosslinking GelMA. **(B)** Workflow showing the approach used to microfabricate chips and to perform subsequent hydrogel patterning. **(C)** Bright-field images showing clearly defined and cell-seedable hydrogel scaffolds. **(D)***(i)* Hydrogel incubated with 4 kDa-TRITC (orange) for 24 hours. Microchannels were filled with 70 kDa-FITC (green) to enhance contrast. *(ii)* yz reconstruction shows square channel geometry with vertical walls 50 µm in height.

We first determined the optimal GelMA prepolymer concentration to prepare a solution suitable for the microfluidic chips. GelMA solutions exhibit a concentration-dependent sol-gel transition between 25-35 °C^57,59,60^. Solutions with concentrations higher than 6% undergo a rapid sol-gel transition upon cooling to room temperature (RT) and are therefore unsuitable for injection into microfluidic channels. GelMA crosslinking at concentrations ranging from 1%-5%, with varying UV doses up to 90 mJ/mm^2^ with 0.1% LAP as the photoinitiator was tested. At 1% GelMA, no visible crosslinking was observed. At 2.5% GelMA, defined structures were observed but they proved fragile and unsuitable for perfusion (data not shown). However, using 5% GelMA, a commonly used concentration, highly defined structures were formed (Figure 1C).

To optimize the LAP concentration, a range of 0.01%-0.5% was tested at UV doses up to 30 mJ/mm^2^. Concentrations lower than 0.05% did not induce GelMA crosslinking, but at 0.1%, well-defined structures were visible even at a UV dose as low as 10 mJ/mm^2^. This corresponds to approximately 4 seconds of exposure per FoV when using a 5X objective. At higher concentrations of LAP, we observed rapid formation of highly defined structures. However, we also detected unwanted crosslinking inside the fluidic channels, which obstructed flow and limited perfusion. This is likely due to long-living radicals that diffuse into the lumens and, at higher concentrations, crosslink the prepolymer^61^. Therefore, a LAP concentration of 0.1% was selected as optimal. This concentration has also been widely reported in the literature.

In summary, GelMA concentrations of 5% and LAP concentrations of 0.1% enabled on-chip patterning of preselected hydrogel structures and scaffolds that are suitable for perfusion and cell seeding (Figure 1C). Hydrogel patterning of a single gel chamber takes 20-30 seconds depending on the required UV-dose (e.g. 20 mJ/mm^2^). Consequently, we were able to generate 20 patterned chips (3 channels per chip) per hour, which is relatively high for hydrogel patterning. In addition, we imaged the shape of the hydrogel using confocal microscopy (Figure 1D). This was achieved by incubating the patterned hydrogels with low molecular weight dextran (4 kDa) and counter-staining the hydrogel channels with high-molecular weight dextran (70 kDa). The cross-sectional view of the 3D reconstruction shows that the hydrogel walls are vertical (Figure 1D). These straight features are possible because the system is based on maskless photolithography and uses collimated light. Similar straight walls from this system have been reported previously^53^. We also observed straight microstructure profiles in the hydrogels when microfabricating SU-8 structures using the system^42^.

### Characterization of patterned GelMA hydrogels

Next, we investigated the controllability of hydrogel parameters. We first tested whether GelMA stiffness can be controlled using different UV doses. The stiffness of GelMA hydrogels depends on the UV dose as well as precursor and photoinitiator concentrations^57,60,62^. Stiffness is important in tissue engineering as there is a natural variability between organs and tissues and it affects the behavior of cells. The elastic modulus of soft-tissue can vary widely, for example, from less than 1 kPa for brain tissue to 1 MPa for intestinal or nerve tissue ^63^.

To mimic the on-chip hydrogel patterning process, we used a “spacer” and removable PDMS “ceiling” (see Materials and Methods). Nanoindentation was used to measure the local mechanical properties of the *in situ*-generated hydrogels, in preference to bulk characterization. The results showed a positive relationship between UV dose and storage modulus (Figure 2A and B). Specifically, the average storage modulus increased from 2.52 kPa with a 10 mJ/mm^2^ UV dose to 4.52 kPa with a 40 mJ/mm^2^ UV dose. Higher UV doses resulted in more crosslinking, leading to an increase in hydrogel stiffness. A plateau was observed at 40 mJ/mm^2^, indicating that continued crosslinking under these conditions would not increase stiffness further, possibly due to depletion of reactive methacrylate groups and/or steric hindrance^57^.

**Figure 2.**
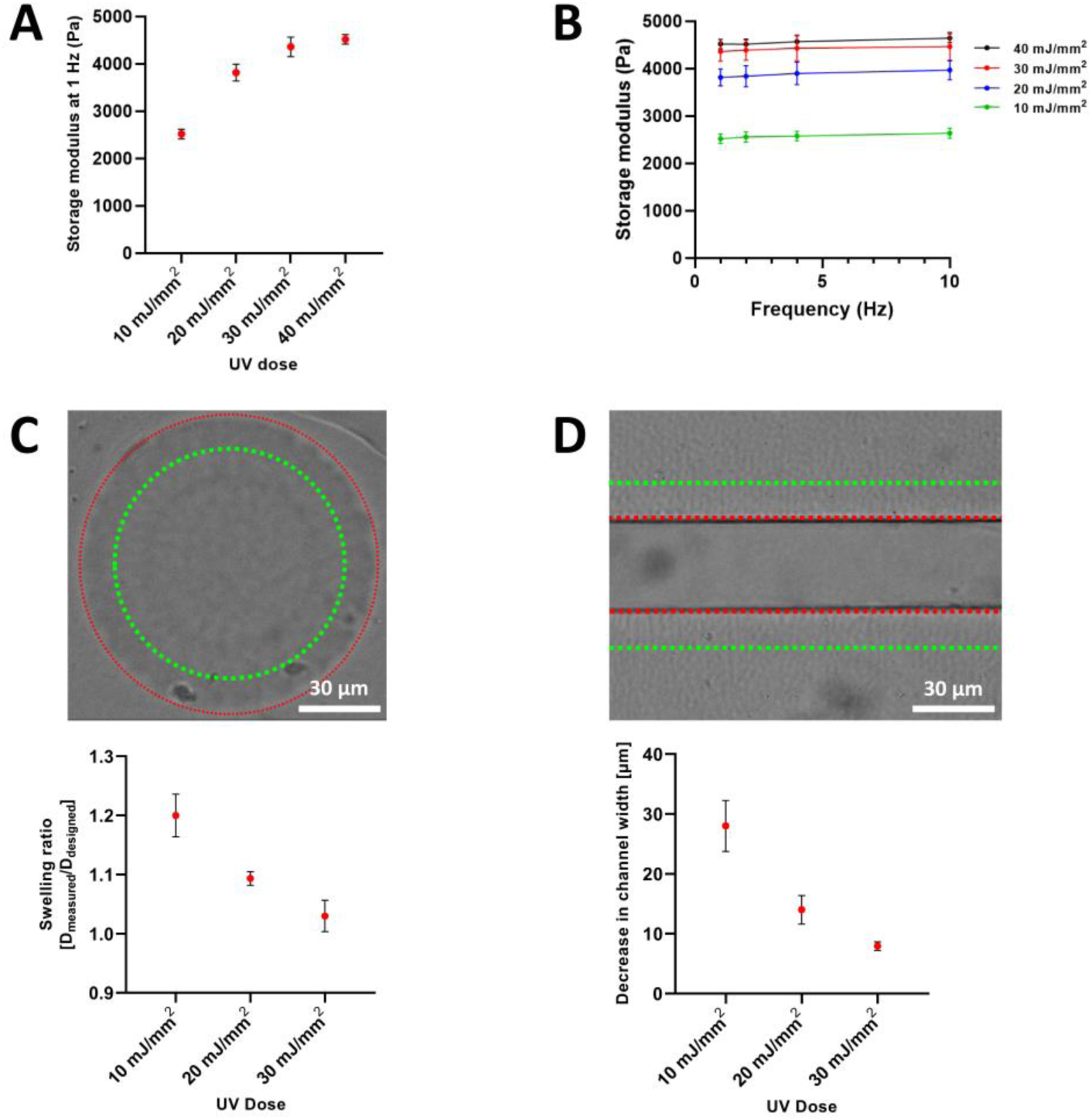
Basic hydrogel characterization. **(A)** Graph showing storage modulus at 1 Hz of patterned GelMA hydrogels crosslinked using increasing laser doses. **(B)** Graph showing storage modulus during 1 Hz – 10 Hz frequency sweep of GelMA hydrogels crosslinked using increasing laser doses. **(C)** Representative bright-field image that was used to determine swelling ratio of 100 µm circular hydrogel structures. The green dotted line shows the designed structure dimensions, while the red dotted line shows the boundary of the actual hydrogel structure after swelling. The graph below displays the quantification of this swelling. **(D)** Representative bright-field image that was used to determine channel narrowing due to hydrogel swelling. The green dotted line shows the designed channel width, while the red dotted line shows the boundary of the hydrogel channel after swelling. The graph below displays the quantification of this narrowing.

The stiffnesses achieved under the conditions tested fell within the range of soft tissue. However, it is important to note that the bottom of the chip is glass and that the top is PDMS, with stiffness several orders of magnitude larger than the hydrogel, making them less physiologically relevant. Nevertheless, since the stiffness and elastic properties of the hydrogel can be tuned, the lateral channel deformation upon increased flow can also be controlled. We observed this effect with increased flow, followed by a return to the relaxed state when flow was stopped (data not shown). Although beyond the scope of this study, modulating the UV laser dose enables control over GelMA stiffness and, consequently, channel compliance. Overall, this offers opportunities to incorporate vascular compliance in engineered microvasculature.

Basic swelling characterization of the hydrogel structures on-chip was then performed using bright-field imaging. Circular structures, 100 µm in diameter, were patterned using UV doses of 10, 20 and 30 mJ/mm^2^, followed by incubation for 24 hours at 37 °C. The final diameters of the swollen structures (Figure 2C, red dotted line) were compared to the photomask dimensions (Figure 2C, green dotted line). Dose-responsive swelling of the hydrogels was observed (Figure 2C, graph), with 10 mJ/mm^2^ resulting in the greatest swelling (ratio of 1.2) and 30 mJ/mm^2^ resulting in the least swelling (ratio of 1.02).

Furthermore, we anticipated narrowing of patterned channels after swelling. To investigate this, the final width of patterned hydrogel channels (Figure 2C, red dotted line) was compared to the photomask dimensions (Figure 2D, green dotted line, 60 µm wide). This revealed an average reduction of 28 µm in channel width when using 10 mJ/mm^2^, a reduction of 14 µm at 20 mJ/mm^2^ and 8 µm at 30 mJ/mm^2^ (Figure 2D, graph).

GelMA swelling is a well-known phenomenon; it is dependent on crosslinking density and the molecular weight between crosslinks, as it affects the water content^57,60^. Therefore, swelling is an important consideration when performing on-chip hydrogel patterning, as it influences the final dimensions of the scaffold.

### Engineering perfusable microvasculature inside microfluidic chips

To engineer microvascular networks inside microfluidic chips, we patterned hydrogel microchannels with a width of 25 µm (Figure 1C). As these scaffolds are readily perfusable, we used passive pumping to populate the scaffolds with hiPSC-ECs.

To assess the coverage of the microchannels by hiPSC-ECs, we used live-cell imaging and confocal microscopy in combination with an alpha-tubulin-GFP hiPSC reporter line. The hiPSC-ECs were thus marked by GFP. Within 2 days or less after seeding, the hiPSC-ECs had covered the channels entirely although they did not follow the square channel geometry as expected. Instead, they formed round and tubular capillary-like structures (Figure 3A). These capillaries exhibited diameters ranging between 20-25 µm. We hypothesize that this may have been caused by the spatial confinement of the ECs within the microchannels^64–66^.

**Figure 3.**
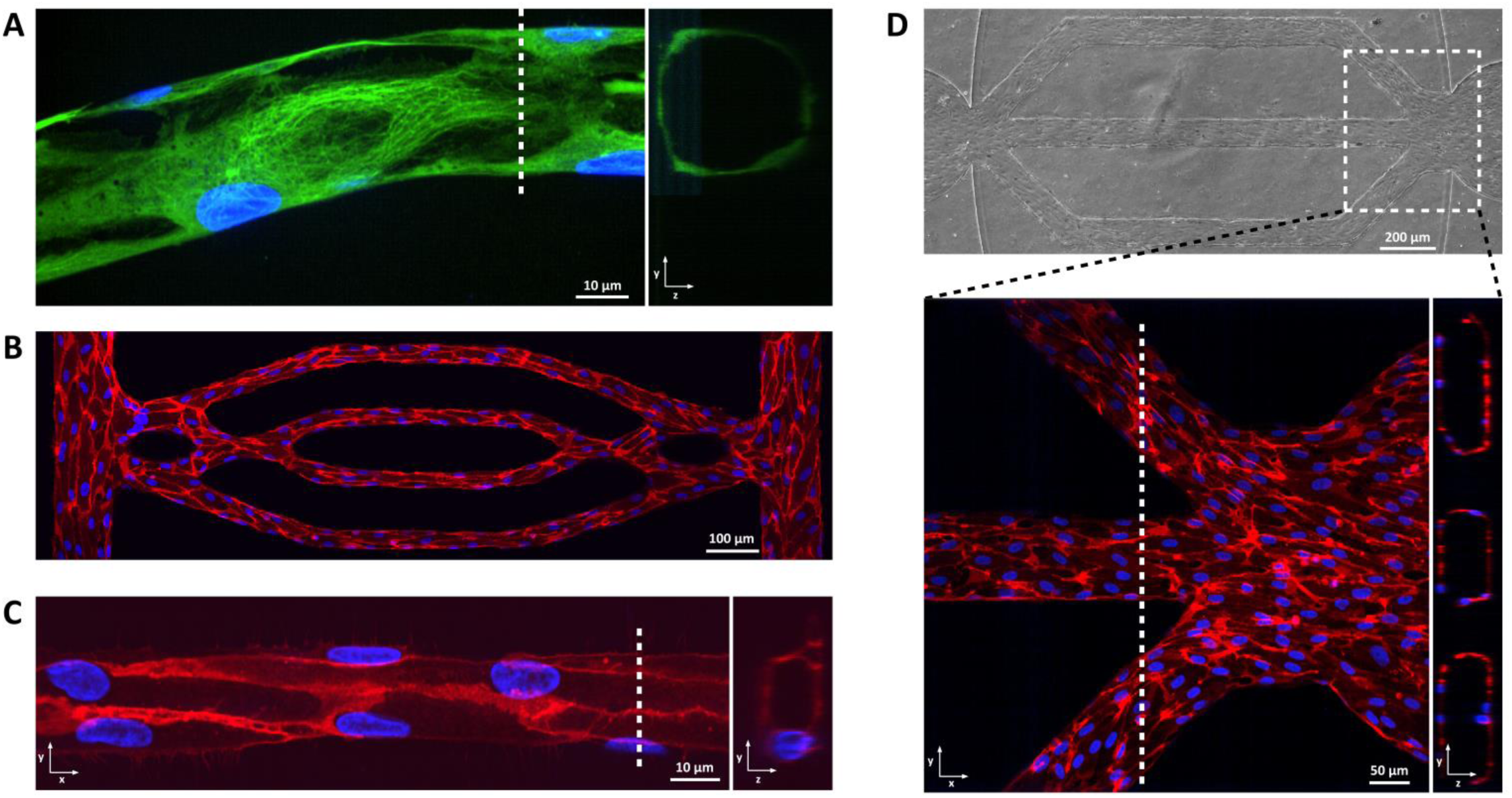
Engineered microvasculature. **(A)** Representative confocal fluorescence image of a section of an engineered microvascular network showing hiPSC-ECs forming a capillary-like structure (green: alpha-tubulin-GFP reporter, blue: hoechst). **(B)** Confocal overview image of an engineered microvascular network (red: CD31, blue: DAPI). **(C)** Representative confocal immunofluorescence image of a capillary-like structure displaying the presence of CD31 (red) near cell-cell junctions. Image displays xy and yz cross-sectional perspectives. **(D)** Bright-field image of engineered arteriole-sized microvasculature 100 µm in width. The inset demonstrates an immunofluorescence image showing CD31 positivity of hiPSC-ECs and their coverage of microchannel lumens. Image displays xy and yz cross-sectional perspectives. White dotted lines indicate cross-sectional planes.

Immunocytochemistry demonstrated clear expression of the pan-EC marker CD31 in engineered microvascular networks (Figure 3B). This suggests that the GelMA hydrogel in its patterned state not only supports cell adhesion and spreading but also maintains EC identity. The networks formed rapidly because we used high cell-density seeding (32*10^6^ cells/mL) whereas in most studies, lower cell densities are used and ECs need to proliferate and migrate extensively to cover the channel walls, or need to invade the hydrogel^33,67^.

Upon closer examination of the capillary-like structures, expression of CD31 was observed specifically near cell-cell junctions (Figure 3C). Furthermore, cross-sectional views demonstrated that capillaries could have diameters as narrow as 10 µm.

To investigate whether larger caliber microvasculature could be generated, we fabricated and seeded scaffolds in channels with 100 µm width (Figure 3D). Within 2 days, the cells were confluent and fully covered the microchannels. In contrast to the capillary-like channels, the ECs followed geometry in these larger channels, as evident in the cross-sectional view (Figure 3D, inset). This is probably because the cells are able to spread entirely across all four planes inside the channels, thereby forming vessels with a square-like geometry.

In summary, the method described here allows facile and rapid engineering of microvasculature having preselected designs with diameters as small as 10 µm.

### Multi-compartment chips for co-culture combinations

To enable co-culture of hiPSC-ECs with other cell types, we introduced compartments into the chip by a simple modification: instead of only one gel inlet on each side of the gel chamber, we added an additional outlet on each side (Figure 4A). By injecting the prepolymer solution via one of the gel inlets and using a photomask that only crosslinks GelMA in the central part of the gel chamber (Figure 4B), we created a 3-compartment chip comprising a central vascular compartment and two side compartments. These side compartments could be flushed with PBS after crosslinking the vascular compartment to remove unreacted GelMA, and subsequently seeded with cells through the additional outlets.

**Figure 4.**
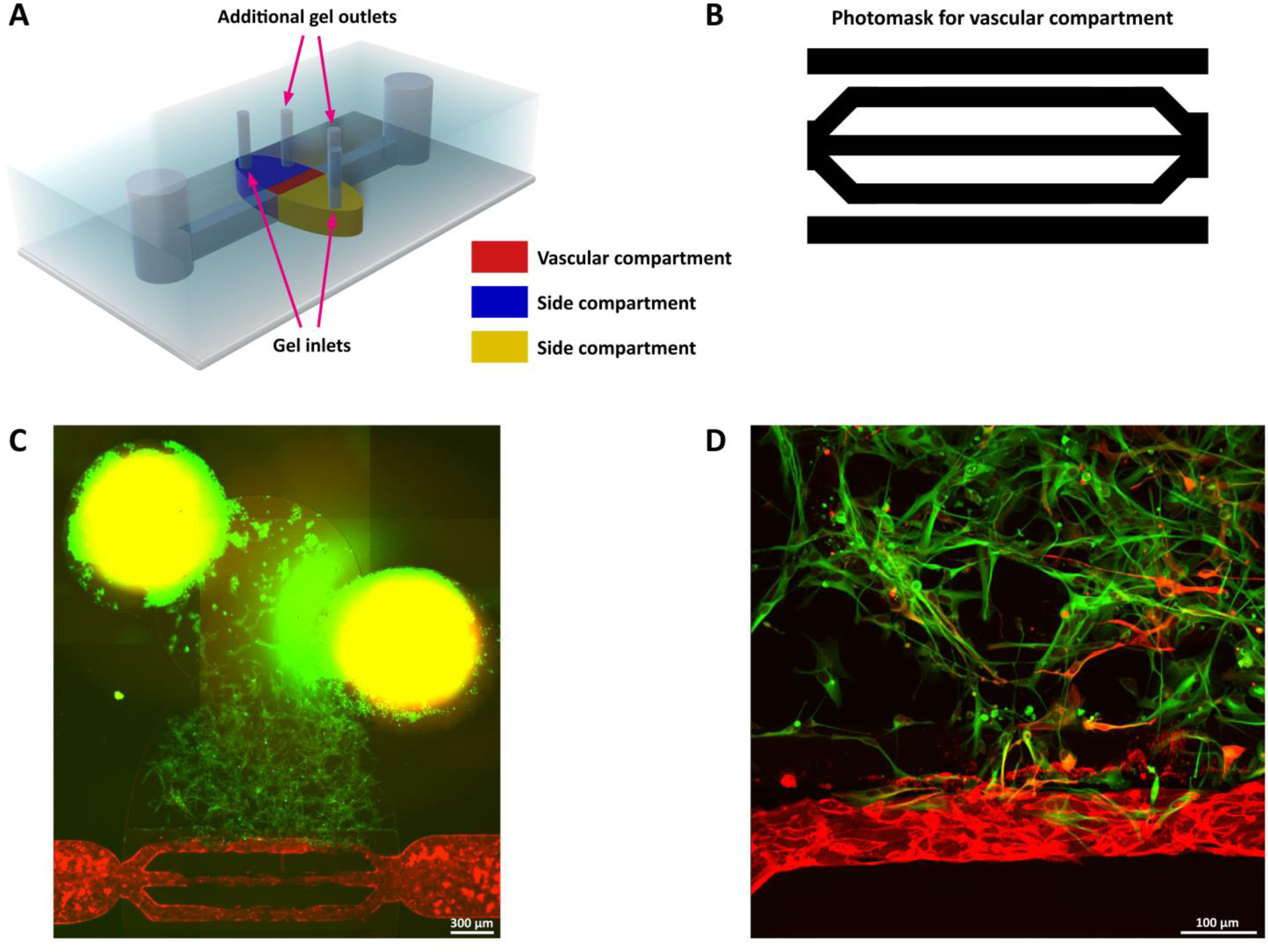
Compartmentalized chip for co-cultures. **(A)** Schematic of a compartmentalized chip generated by patterning a central vascular compartment, thereby dividing the gel chamber into three different compartments. The additional gel outlets enable: (1) flushing of unpolymerized GelMA solution, (2) injection, and (3) ejection and removal of excess fibrin gel-cell suspension. **(B)** Digital photomask used to pattern the central vascular compartment with 100 µm wide microchannels. White is exposure, black is no exposure. **(C)** Overview image of a central vascular compartment populated with hiPSC-ECs and an adjacent compartment containing a fibrin hydrogel with hiPSC-astrocytes. The gel inlet and additional outlet are clearly visible due to a bright green signal caused by the abundant presence of hiPSC-astrocytes. **(D)** Confocal image showing a section of a central vascular compartment and an adjacent compartment. hiPSC-astrocytes are interacting with and migrating towards hiPSC-ECs in the central vascular compartment (day 3). Red: CD31, green: hiPSC-astrocytes with alpha-tubulin-GFP reporter.

As a proof of concept for a relevant co-culture, hiPSC-astrocytes were resuspended in a fibrin hydrogel and carefully injected into one side compartment and hiPSC-ECs cultured in the vascular compartment as earlier. After 3 days, hiPSC-ECs lined the microchannels, while the hiPSC-astrocytes had spread out and were migrating towards the vascular compartment (Figure 4C). The images demonstrated that the GelMA could withstand the injection of the cell suspension and remain in place. Confocal imaging revealed interactions between the astrocytes and the vascular compartment (Figure 4D). This method of compartmentalization is thus feasible for co-culture of hiPSC-ECs with one or more additional cell types.

This relatively straightforward modification of the microfluidic chips means that the blood-brain barrier for example could be modeled, a structure of significant interest in biomedical research. Other examples include modeling the lymphatic vasculature in combination with different tissue types to study lymphatic drainage and immune cells.

### Patterning functional valves

Facile patterning of microvascular scaffolds enabled us to explore other aspects of vascular engineering. Specifically, we patterned valve leaflets (Figure 5A(i)) inside 100 µm wide hydrogel channels. First, the microchannels were patterned as before and the valves were then immediately patterned inside these channels using a UV dose of 20 mJ/mm^2^. The photomask shown in Figure 5A(ii) was designed to resemble the shape of venous valves. They were capable of opening when flow was applied (Figure 5A(i), top image) and closing when flow was stopped (Figure 5A(i), bottom image). This functionality indicated that the valves could physically move and regulate the opening and closing of the channels. Additionally, the valves demonstrated elastic deformation upon flow reversal (Figure 5A(iii)).

**Figure 5.**
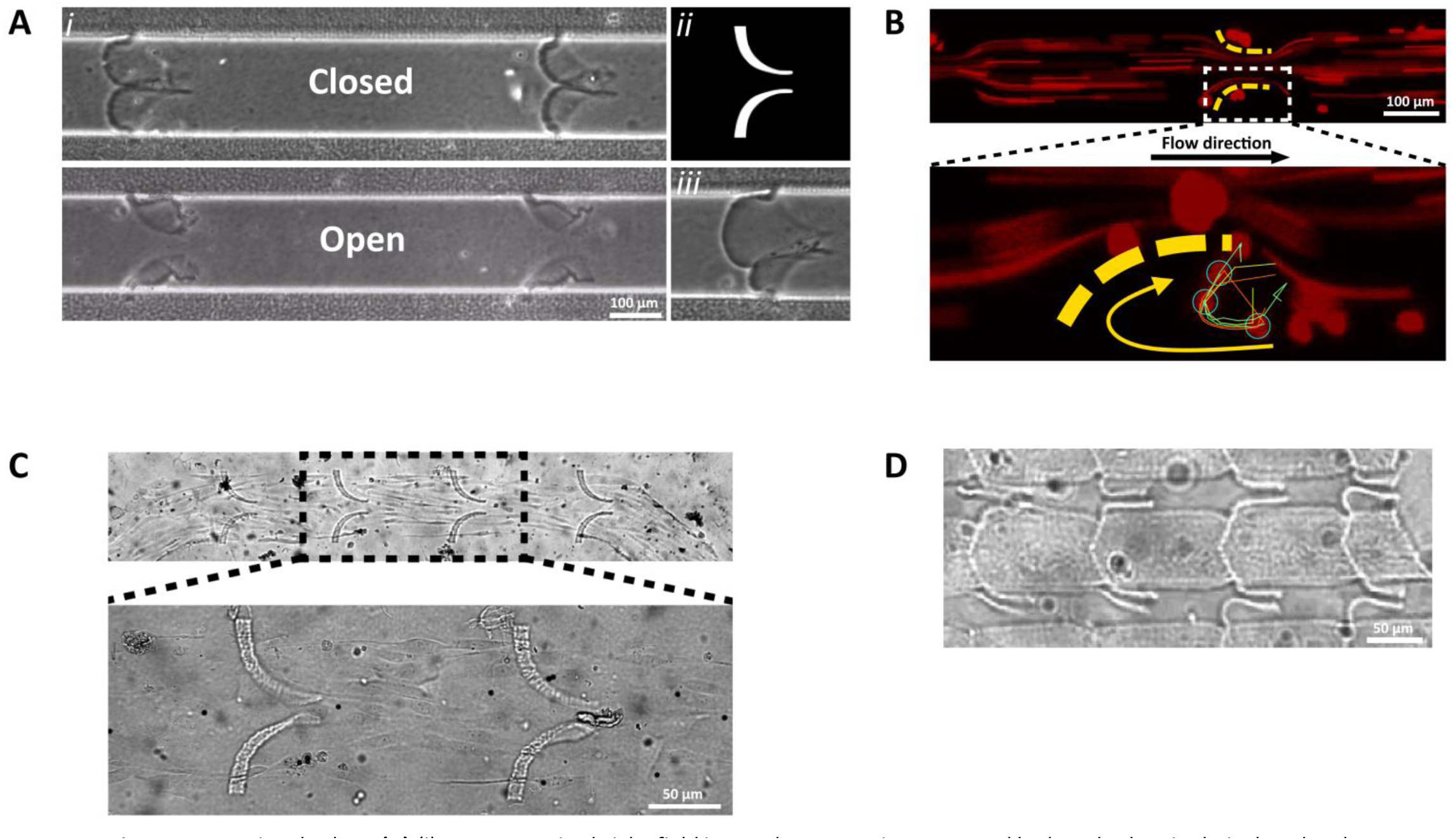
Functional valves. **(A)** *(i)* Representative bright-field image demonstrating patterned hydrogel valves in their closed and open state inside hydrogel microchannels 100 µm in width. *(ii)* Digital photomask used to pattern valves inside the hydrogel microchannels. *(iii)* Representative bright-field image of elastically deformed valves upon flow reversal. **(B)** Single image of a sequence of recorded fluorescence images indicating beads flowing past valves (yellow dotted lines). The inset visualizes vortical flow near a valve tip, analyzed using single particle tracking. The colored lines indicate tracks of beads that underwent vortical flow during recording. Beads that were undergoing vortical flow in this frame are indicated by cyan circles. **(C)** Bright-field image of hydrogel valves inside a live microvessel (containing hiPSC-ECs) that was engineered before valve patterning. **(D)** Representative bright-field image of hydrogel valves patterned inside microchannels that are 50 µm in width.

Modeling valves and valve-related hemodynamic processes *in vitro* has been demonstrated previously using microfluidic chips. Such chips have proven valuable for evaluating the effects of channel geometry and flow dynamics on particle behavior (e.g. beads, platelets or red blood cells) and could thus offer insight into the pathophysiology of diseases such as thrombosis^68–73^.

Next, we examined the presence of vortical flow near the valve tips by manually adding fluorescent beads to flow through the channels. Vortical flow that occurs near venous valve tips *in vivo*, has also been studied *in vitro* using hydrogel valves^70–72^. These studies characterized flow dynamics around the valves, which are critical drivers of thrombosis-related processes. Laminar fluid flow was also visualized by the motion blur of the beads (Figure 5B(i)). Through single particle tracking near the valve tips, we were able to detect vortical flow from the particle tracks shown in Figure 5B(ii). In this image, a single frame depicts colored particle tracks indicating the trajectories (curved yellow arrow) of several particles identified during the recorded frames near a valve tip (valve indicated by the yellow dotted line). Three particles identified undergoing vortical flow are highlighted by cyan-colored circles in this frame.

To assess whether valves could be patterned inside live endothelialized microchannels, we first generated microvessels as described earlier. Subsequently, the GelMA prepolymer solution was reintroduced, and the valves were patterned. Figure 5C shows the resulting microvessel with the valves present. The formation of functional microvasculature containing valves was thus demonstrated.

Finally, we successfully patterned functional valves inside microchannels that were 50 µm wide. Despite being small, these valves retained the ability to open and close the channels (Figure 5D). To our knowledge, functional hydrogel valves at this scale have not been generated previously. Importantly, such small valves resemble microscopic venous valves (MVVs) and lymphatic vessels, enabling conditions that affect these valves to be modelled. MVVs have been identified in the human body, with evidence suggesting their abundance in microvasculature measuring less than 100 µm and as small as 20 µm^74,75^.

Whilst there have been other reports of these types of valves being modelled in different chip designs, we showed here that valves of multiple sizes and/or shapes could be generated inside microvascular hydrogel scaffolds using a single type of system. These valves exhibited proper opening and closing functionality, demonstrating flexibility and elastic deformation. Moreover, their design is adaptable and can be optimized for other applications than those described here. For instance, changes in their shape or hydrogel stiffness, could allow the impact on vortical flow formation and hemodynamics to be determined^70^.

## Limitations

The method we developed facilitates the microfabrication of chips and subsequent patterning of microvascular scaffolds with relative ease, hydrogel patterning is straightforward and many chips per hour can be produced, but also has limitations. Since the method is based on photolithography, which means that there is complete crosslinking of everything in the light path and that it is rapid, there is limited control over microstructure height. However, hydrogel height control inside enclosed chambers has been achieved using grayscale projections, oxygen control, and a benzophenone-based photoinitiator^53^. Using our approach, the PRIMO system could control the height of crosslinked hydrogel structures. Nevertheless, this does not match the exceptional control of 2PP or 3D printing. As a result, geometries achievable are limited primarily to square shapes. Additionally, the glass bottom and PDMS top of the chips have a significantly higher stiffness compared to the more physiologically relevant hydrogel-based walls. These differences may affect the behavior of cells cultured on the top or bottom of the chip. Finally, cell encapsulation presents challenges due to the phototoxic nature of the patterning process. The exposure wavelength (375 nm) combined with the photoinitiator LAP can induce cytotoxicity by generating free radicals during the patterning process^23^.

## Conclusion

We developed here a straightforward method for engineering functional and perfusable microvasculature with predetermined designs inside custom microfabricated chips using a single DMD-based setup for maskless photolithography (PRIMO). Cell-receptive GelMA hydrogel scaffolds with controllable and *in vivo*-like mechanical properties could be patterned, simply by adjusting the UV laser dose. We successfully engineered microvascular networks containing capillary-like tubular structures as small as 10 µm, as well as larger arteriole-sized microvessels. Moreover, a simple modification of the chips allowed compartmentalization, enabling co-culture of microvasculature with other cell types such as hiPSC-astrocytes. Additionally, we demonstrated the patterning of functional valves inside hydrogel microchannels, capable of opening and closing the channels, even with live microvessels. This opens avenues for studying valve-related hemodynamics *in vitro* and disease modeling such as thrombosis. Overall, the precise level of control demonstrated in this study enables the rapid engineering of a diverse array of microvasculature on-chip.

## Materials and Methods

### Maskless photolithography to generate glass master mold

Glass substrates containing SU-8 microstructures were used as master molds and were fabricated as previously described^42^. Briefly, glass substrates spin coated with a 50 μm thick layer of SU-8-2075 (Kayaku Advanced Materials, Inc) were placed into an adjustable microscope holder for backside UV exposure. A maskless DMD-based photolithography system (PRIMO, Alvéole) connected to a Leica DMi8 inverted microscope with motorized stage was used for exposure. Binary digital photomasks were designed in an open-source vector graphics editor (Inkscape). The digital photomasks were then loaded into the Leonardo software (Alvéole) and projected by the system onto the substrate via a 5X/0.15NA objective using a 6 mJ/mm^2^ laser dose. Finally, post-exposure bake and subsequent development of SU-8 were performed as described earlier.

### Mold replication

The glass master mold was replicated by first generating a master copy followed by replication using an epoxy-resin to make durable copies. The microfabricated glass SU-8 master mold was mounted in a 60 mm diameter polystyrene Petri dish (Greiner Bio-One, Germany) using a thin layer of Polydimethylsiloxane (PDMS, Sylgard 184, DOW Chemicals, USA), after which a master copy was made using Smooth-Sil 940 (Smooth-On, USA) according to manufacturer’s instructions. Smooth-Sil 940 components were mixed and degassed for 30 min at RT under high vacuum. 10 g was then poured into the Petri dish and cured at RT for 24 hours. The Smooth-Sil 940 was subsequently demolded from the Petri dish by carefully cutting a 35 mm circle around the microstructures using a scalpel. This procedure was repeated five times afterwards (using the glass master mold). Next, the six master replicates were mounted in a polypropylene box and Epoxacast 670 HT (Smooth-On) was casted on top. Epoxacast components A and B were first mixed according to manufacturer’s instructions and subsequently mixed with thinner (Epic Thinner, Smooth-On) to reduce resin viscosity. The mixture was then degassed at RT for 30 min under high vacuum. The cast was cured at RT for 12 hours followed by 3 hours at 80 °C and 24 hours at 70 °C. The resulting epoxy molds were then used for soft lithography to fabricate PDMS chips.

### Soft lithography and plasma bonding

PDMS was mixed in a 10:1 (base:curing agent) mass ratio and degassed for 30 min at RT. Five grams of PDMS were then poured into the epoxy molds and placed at 70 °C for 24 hours. After curing, PDMS was allowed to cool to RT after which the PDMS was gently peeled off. The inlets of the fluidic channels were punched using a 3 mm biopsy puncher. For the gel chamber outlets and inlets, a 1.2 mm biopsy puncher was used. Finally, PDMS chips were bonded to 35 mm glass coverslips (Menzel-Glaser, #1) using air plasma (50 W, 50 KHz) for 1 min (Cute, Femto Science Inc., Korea).

### Hydrogel scaffold and valve patterning

62.5 mg/mL gelatin methacrylate (GelMA, 300 Bloom, degree of substitution 60%, 900622, Sigma-Aldrich) was dissolved at 37 °C in phosphate-buffered saline (PBS) (w/v), aliquoted and stored at 4 °C. Before using, the solution was pre-warmed to 40 °C and mixed 4:1 with freshly prepared 5 mg/mL lithium phenyl-2,4,6 trimethylbenzoylphosphinate (LAP, Sigma-Aldrich) in PBS (w/v) to obtain a hydrogel working solution containing 5% GelMA and 0.1% LAP. Finally, the working solution was prewarmed to 40 °C and 8 µL was then injected into the gel chambers of the microfluidic chips. The filled microfluidic chips were immediately placed into an adjustable microscope holder for exposure using the DMD-based system (PRIMO, Alvéole). A digital photomask containing the required GelMA pattern was designed in Inkscape, loaded into Leonardo (Alvéole) and projected by the system onto the gel chamber via a 5X/0.15NA objective using a 20 mJ/mm^2^ laser dose. Microfluidic chips were then transferred to a hotplate set at 40 °C and flushed for 15 minutes using prewarmed PBS to remove un-crosslinked GelMA and excess photoinitiator. When valves were required, digital photomasks containing valves were loaded and valves were then patterned (using a 20 mJ/mm^2^ laser dose) inside the patterned hydrogel microchannels prior to flushing.

To improve cell adhesion, chips were incubated with a fibronectin in distilled water solution (50 µg/mL, bovine, Thermo Fisher Scientific) for 2 h at RT or overnight at 4 °C. Chips were flushed and pre-filled with EC culture medium (EGM-2) before usage.

### Determination of hydrogel swelling ratio

A digital photomask containing circular patterns of 100 µm in diameter was designed in Inkscape and projected onto the chip gel chamber using the PRIMO system at laser doses of 10, 20 and 30 mJ/mm^2^ using the Leonardo software (Alvéole) as described above. Microfluidic chips were flushed with EGM-2 and placed inside an incubator at 37 °C for 24 hours. After incubation, the structured hydrogels inside the chip were imaged using an EVOS M7000 fluorescence microscope (ThermoFisher Scientific) and compared to the dimensions of the digital photomask. Similarly, straight hydrogel channels 60 µm in width were patterned and flushed as described above. After 24h incubation, the hydrogel channels were imaged and compared with the dimensions of the digital photomask.

### Hydrogel stiffness (nanoindentation)

A nanoindenter (Chiaro, Optics11Life, The Netherlands) was used to characterize the crosslinked GelMA hydrogels mechanically. To perform nanoindentation, the on-chip hydrogel patterning process was mimicked. Briefly, a 50 µm high spacer was placed onto a glass coverslip after which 20 uL of pre-warmed hydrogel solution was pipetted in the middle. A block of PDMS was then placed on top so that the solution was confined between the glass, spacer and the PDMS. This 50 µm high solution was then crosslinked using the PRIMO system with the specified laser dose. Afterwards, the coverslip with hydrogel and PDMS was transferred to a hotplate set at 40 °C and prewarmed PBS was pipetted between the glass and the PDMS. After 10 minutes, the PDMS was carefully removed and the patterned hydrogel was then accessible for nanoindentation. An indentation probe with 0.17 N/m cantilever stiffness and a spherical tip of 53.5 µm radius was used (Optics11Life). Dynamic mechanical analysis was performed using a 1 Hz–10 Hz frequency sweep and a 4 µm indentation depth.

### Cell culture

The following hiPSC lines were used: LUMC0020iCTRL (generated from skin fibroblasts, https://hpscreg.eu/cell-line/LUMCi028-A) and The Allen Cell Collection line AICS-0012 (generated from skin fibroblasts, https://hpscreg.eu/cell-line/UCSFi001-A-2) with mEGFP insertion site at TUBA1B. hiPSCs for generation of hiPSC-ECs were maintained on recombinant vitronectin-coated plates in TeSR-E8 medium (05940, StemCell Technologies, Canada) and passaged once a week using Gentle Cell Dissociation Reagent (07174, StemCell Technologies). hiPSCs for generation of hiPSC-astrocytes were cultured on Matrigel-coated (354230, BD Biosciences) plates in mTeSR1 medium (05850, StemCell Technologies) and mechanically passaged once a week using dispase solution 1 mg/mL (17105-041, Gibco). hiPSC-ECs were derived and maintained as previously described^76,77^. hiPSC-astrocytes were derived and maintained as previously described^78^.

### Seeding of scaffolds with hiPSC-ECs

hiPSC-ECs were dissociated using TrypLE™ (ThermoFisher Scientific) and resuspended at a concentration of 32*10^6^ cells/mL in EGM-2 supplemented with PenStrep (25 Units/mL). 5 µl of cell suspension was carefully pipetted onto the bottom of the inlet using a P10 pipette tip causing the cells to be passively pumped inside the scaffold. After 5 seconds, 5 µl of cell suspension was also pipetted on the bottom of the outlet. Subsequently, after 5 seconds, the cell suspension was retrieved from the inlet and outlet using a P10 pipette. As such, flow was halted and cells were then allowed to attach for 30 minutes at 37 °C in a humidified polypropylene box. 40 µl of pre-warmed EGM-2 was added to the inlets and outlets and chips were then incubated for 2 hours at 37 °C. Afterwards, chips were transferred to a rocking platform (OrganoFlow L, MIMETAS B.V., Leiden, The Netherlands) and cultured for at least 48 hours under bidirectional flow (7-degree inclination at 8 min intervals) in a humidified polypropylene box. Medium was refreshed every day by aspiration followed by addition of 40 µL to the inlets and outlets.

### hiPSC-astrocyte co-culture

To introduce compartmentalization, an additional gel outlet was punched on each side of the gel chamber before chip assembly. After assembly, the chip was filled with 8 µL of GelMA solution as described earlier. An adapted digital photomask was designed where only the middle section of the gel chamber is exposed during patterning, thereby creating a central vascular compartment with two empty side-compartments after flushing. Patterning the hydrogel microchannels of the central vascular compartment was performed as described earlier. After patterning, microfluidic chips were transferred to a hotplate set at 40 °C and the vascular compartment was flushed for 15 minutes using prewarmed PBS. To flush the side-compartments, 10 µL of prewarmed PBS was carefully injected into the gel inlets using a P10 pipette. This resulted in the un-crosslinked GelMA, excess photoinitiator and PBS exiting the chip via the additional gel outlet. This was repeated three times in order to fully eliminate the prepolymer solution from the compartments.

hiPSC-astrocytes were then dissociated using TrypLE™ (ThermoFisher Scientific) and resuspended in EGM-2 medium supplemented with thrombin (4U/mL, T4648, Sigma) at 50*10^6^ cells/mL. Prior to injection into a chip, 5 µL of cell suspension was mixed with 5 µL of fibrinogen solution (6 mg/mL, final concentration 3 mg/mL, 8630, Sigma) by pipetting up and down three times. The resulting astrocyte suspension then contained 25*10^6^ cells/mL and was immediately injected into the gel inlet, thereby filling the side-compartment and displacing the PBS. The excess of the cell suspension could exit the side compartment via the additional gel outlet and was discarded. The chip was subsequently incubated at RT for 15 minutes until the fibrin-gel was polymerized. hiPSC-ECs were then seeded and chips were cultured using EGM-2 medium as described earlier.

### Single-particle tracking

Chips containing patterned hydrogel microchannels with valves inside were fabricated as described above. PTFE tubing (1/16″ OD X 1/32″ ID*, LVF-KTU-15, Elveflow) was connected to the chip inlet (via a 14G blunt-end needle with Luer lock tip) and to a 15 mL syringe.

The syringe was filled with a PBS solution containing 2 µm-sized fluorescent beads (25*10^6^ beads/mL, ThermoFisher Scientific). The solution was then manually flowed through the chip in the proper direction by depressing the syringe plunger, such that the valves in the microchannel open up. Concurrently, to capture bead movement, a region of interest containing valves was recorded at 28 frames/s using an EVOS M7000 microscope (Thermo Fisher Scientific) using a 10X/0.30NA objective. To perform single-particle tracking and to visualize vortical bead tracks, TrackMate (ImageJ plugin) was used^79,80^.

### Immunocytochemistry

All steps below were performed by adding 30 µL of the specified solutions in the inlets and 20 µL in the outlets. Samples were fixed by adding 4% paraformaldehyde (PFA) solution followed by a 15 min incubation at RT. Samples were then washed three times with PBS and permeabilized for 10 min at RT using 0.1% Triton X-100 in PBS (-). Subsequently, the samples were blocked for 1 h at RT using 1% bovine serum albumin (BSA) in PBS (-). Primary antibodies were diluted 1:200 in 1% BSA in PBS(-) and incubated overnight at 4 °C. The primary antibody used was against CD31 (PECAM1, mouse, M0823, Agilent Dako, Santa Clara, CA, USA) to stain ECs. Samples were washed 3 times with PBS (-) afterwards. Secondary antibody (A-21203, Thermo Fisher Scientific) was then diluted 1:300 in 1% BSA in PBS (-) and incubated at RT for 2 h. Cell nuclei were stained using DAPI. Samples were stored in PBS (-) in the dark at 4 °C until imaging.

### Immunofluorescence and live cell imaging

Cells and gels were imaged using a Leica SP8 microscope connected to a Dragonfly 500 spinning disk confocal system (Andor Technology Ltd.) using a 20X, 40X or 63X objective. Post-processing was performed using Imaris 9.5 software (Bitplane, Oxford Instruments). Immunofluorescence and bright-field overview scans were acquired using an EVOS M7000 microscope (Thermo Fisher Scientific) with a 10X/0.30NA objective.

## Acknowledgements

This work was supported by the LymphChip project with project number NWA-ORC 2019 1292.19.019 of the NWA research program ‘‘Research on Routes by Consortia” (ORC), which is funded by the Netherlands Organization for Scientific Research (NWO); the Netherlands Organ-on-Chip Initiative, which is an NWO Gravitation project (024.003.001) funded by the Ministry of Education, Culture and Science of the government of the Netherlands, and The Novo Nordisk Foundation Center for Stem Cell Medicine that is supported by a Novo Nordisk Foundation grant (NNF21CC0073729).

## Notes

### Competing Interest Statement

The authors have declared no competing interest.

## References

1. Glassman, P. M. et al. Targeting drug delivery in the vascular system: Focus on endothelium. Adv Drug Deliv Rev 157, 96–117 (2020).

2. Gutterman, D. D. et al. The Human Microcirculation. Circ Res 118, 157–172 (2016).

3. Reiterer, M. & Branco, C. M. Endothelial cells and organ function: applications and implications of understanding unique and reciprocal remodelling. FEBS Journal 287, 1088–1100 (2020).

4. Osaki, T., Sivathanu, V. & Kamm, R. D. Vascularized microfluidic organ-chips for drug screening, disease models and tissue engineering. Curr Opin Biotechnol 52, 116–123 (2018).

5. Huh, D. et al. Reconstituting Organ-Level Lung Functions on a Chip. Science (1979) 328, 1662– 1668 (2010).

6. Dellaquila, A., Le Bao, C., Letourneur, D. & Simon-Yarza, T. In Vitro Strategies to Vascularize 3D Physiologically Relevant Models. Advanced Science 8, (2021).

7. Tronolone, J. J. & Jain, A. Engineering New Microvascular Networks On-Chip: Ingredients, Assembly, and Best Practices. Adv Funct Mater 31, (2021).

8. Cochrane, A. et al. Advanced in vitro models of vascular biology: Human induced pluripotent stem cells and organ-on-chip technology. Adv Drug Deliv Rev (2018) doi:10.1016/j.addr.2018.06.007.

9. Bhatia, S. N. & Ingber, D. E. Microfluidic organs-on-chips. Nat Biotechnol 32, 760–772 (2014).

10. Ingber, D. E. Human organs-on-chips for disease modelling, drug development and personalized medicine. Nat Rev Genet 0123456789, (2022).

11. Myers, D. R. & Lam, W. A. Vascularized Microfluidics and Their Untapped Potential for Discovery in Diseases of the Microvasculature. (2021) doi:10.1146/annurev-bioeng-091520.

12. Jeon, J. S. et al. Generation of 3D functional microvascular networks with human mesenchymal stem cells in microfluidic systems. Integrative Biology (United Kingdom) 6, 555–563 (2014).

13. Vila Cuenca, M., et al. Engineered 3D vessel-on-chip using hiPSC-derived endothelial- and vascular smooth muscle cells. Stem Cell Reports 16, 2159–2168 (2021).

14. Kim, S., Lee, H., Chung, M. & Jeon, N. L. Engineering of functional, perfusable 3D microvascular networks on a chip. Lab Chip 13, 1489–1500 (2013).

15. Campisi, M. et al. 3D self-organized microvascular model of the human blood-brain barrier with endothelial cells, pericytes and astrocytes. Biomaterials 180, 117–129 (2018).

16. van Duinen, V. et al. Perfused 3D angiogenic sprouting in a high-throughput in vitro platform. Angiogenesis 0, 0 (2018).

17. Kačarević, Ž. P., et al. An Introduction to 3D Bioprinting: Possibilities, Challenges and Future Aspects. Materials 11, 2199 (2018).

18. Miri, A. K. et al. Bioprinters for organs-on-chips. Biofabrication 11, 042002 (2019).

19. Yu, F. & Choudhury, D. Microfluidic bioprinting for organ-on-a-chip models. Drug Discov Today 24, 1248–1257 (2019).

20. Homan, K. A. et al. Bioprinting of 3D Convoluted Renal Proximal Tubules on Perfusable Chips. Sci Rep 6, (2016).

21. Kim, W. et al. 3D Inkjet-Bioprinted Lung-on-a-Chip. ACS Biomater Sci Eng 9, 2806–2815 (2023).

22. Kolesky, D. B. et al. 3D bioprinting of vascularized, heterogeneous cell-laden tissue constructs. Advanced Materials 26, 3124–3130 (2014).

23. Lim, K. S. et al. Fundamentals and Applications of Photo-Cross-Linking in Bioprinting. Chem Rev 120, 10662–10694 (2020).

24. Nguyen, A. K. & Narayan, R. J. Two-photon polymerization for biological applications. Materials Today 20, 314–322 (2017).

25. Faraji Rad, Z., Prewett, P. D. & Davies, G. J. High-resolution two-photon polymerization: the most versatile technique for the fabrication of microneedle arrays. Microsyst Nanoeng 7, 71 (2021).

26. Limongi, T. et al. Three-dimensionally two-photon lithography realized vascular grafts. Biomedical Materials (Bristol) 16, (2021).

27. Brandenberg, N. & Lutolf, M. P. In Situ Patterning of Microfluidic Networks in 3D Cell-Laden Hydrogels. Advanced Materials 28, 7450–7456 (2016).

28. Enrico, A. et al. 3D Microvascularized Tissue Models by Laser-Based Cavitation Molding of Collagen. Advanced Materials 34, (2022).

29. Arakawa, C. et al. Biophysical and biomolecular interactions of malaria-infected erythrocytes in engineered human capillaries. Sci Adv 6, (2020).

30. Rayner, S. G. et al. Multiphoton-Guided Creation of Complex Organ-Specific Microvasculature. Adv Healthc Mater 10, (2021).

31. Pradhan, S., Keller, K. A., Sperduto, J. L. & Slater, J. H. Fundamentals of Laser-Based Hydrogel Degradation and Applications in Cell and Tissue Engineering. Adv Healthc Mater 6, (2017).

32. Cantoni, F., Barbe, L., Pohlit, H. & Tenje, M. A Perfusable Multi-Hydrogel Vasculature On-Chip Engineered by 2-Photon 3D Printing and Scaffold Molding to Improve Microfabrication Fidelity in Hydrogels. Adv Mater Technol 9, (2024).

33. Zhao, N. et al. Engineering the Human Blood–Brain Barrier at the Capillary Scale using a Double-Templating Technique. Adv Funct Mater 32, (2022).

34. Polacheck, W. J., Kutys, M. L., Tefft, J. B. & Chen, C. S. Microfabricated blood vessels for modeling the vascular transport barrier. Nat Protoc 14, 1425–1454 (2019).

35. Bertassoni, L. E. et al. Hydrogel bioprinted microchannel networks for vascularization of tissue engineering constructs. Lab Chip 14, 2202–2211 (2014).

36. Jiménez-Torres, J. A., Peery, S. L., Sung, K. E. & Beebe, D. J. LumeNEXT: A Practical Method to Pattern Luminal Structures in ECM Gels. Adv Healthc Mater 5, 198–204 (2016).

37. Chrobak, K. M., Potter, D. R. & Tien, J. Formation of perfused, functional microvascular tubes in vitro. Microvasc Res 71, 185–196 (2006).

38. Dudley, D., Duncan, W. M. & Slaughter, J. Emerging digital micromirror device (DMD) applications. MOEMS Display and Imaging Systems 4985, 14 (2003).

39. Takahashi, K. & Setoyama, J. A UV-exposure system using DMD. Electronics and Communications in Japan (Part II: Electronics) 83, 56–58 (2000).

40. Li, X. et al. Plasmonic nanohole array biosensor for label-free and real-time analysis of live cell secretion. Lab Chip 17, 2208–2217 (2017).

41. Zhang, Y. et al. User-defined microstructures array fabricated by DMD based multistep lithography with dose modulation. Opt Express 27, 31956 (2019).

42. Kasi, D. G. et al. Rapid Prototyping of Organ-on-a-Chip Devices Using Maskless Photolithography. Micromachines (Basel) 13, 49 (2021).

43. Souquet, B., Opitz, M., Vianay, B., Brunet, S. & Théry, M. Manufacturing a Bone Marrow-On-A-Chip Using Maskless Photolithography. in Bone Marrow Environment: Methods and Protocols (eds. Espéli, M. & Balabanian, K.) 263–278 (Springer US, New York, NY, 2021). doi:10.1007/978-1-0716-1425-9_20.

44. Blin, G. Quantitative developmental biology *in vitro* using micropatterning. Development 148, (2021).

45. Jung, Y., Lee, H., Park, T. J., Kim, S. & Kwon, S. Programmable gradational micropatterning of functional materials using maskless lithography controlling absorption. Sci Rep 5, (2015).

46. Xiong, Z., Kunwar, P. & Soman, P. Hydrogel-Based Diffractive Optical Elements (hDOEs) Using Rapid Digital Photopatterning. Adv Opt Mater 9, (2021).

47. Oliver, C. R., Gourgou, E., Bazopoulou, D., Chronis, N. & Hart, A. J. On-demand isolation and manipulation of C. elegans by in vitro maskless photopatterning. PLoS One 11, (2016).

48. Carbonell, C. et al. Polymer brush hypersurface photolithography. Nat Commun 11, (2020).

49. Yang, W., Yu, H., Liang, W., Wang, Y. & Liu, L. Rapid fabrication of hydrogel microstructures using UV-induced projection printing. Micromachines (Basel) 6, 1903–1913 (2015).

50. Mitmoen, M. & Kedem, O. UV-and Visible-Light Photopatterning of Molecular Gradients Using the Thiol-yne Click Reaction. ACS Appl Mater Interfaces 14, 32696–32705 (2022).

51. Lowry Curley, J., Jennings, S. R. & Moore, M. J. Fabrication of micropatterned hydrogels for neural culture systems using dynamic mask projection photolithography. Journal of Visualized Experiments (2011) doi:10.3791/2636.

52. Strale, P. O. et al. Multiprotein Printing by Light-Induced Molecular Adsorption. Advanced Materials 28, 2024–2029 (2016).

53. Pasturel, A., Strale, P. O. & Studer, V. Tailoring Common Hydrogels into 3D Cell Culture Templates. Adv Healthc Mater 9, (2020).

54. Mazari-Arrighi, E. et al. Construction of functional biliary epithelial branched networks with predefined geometry using digital light stereolithography. Biomaterials 279, 121207 (2021).

55. Robineau, P., Béal, J., Pons, T., Jaffiol, R. & Vézy, C. Micropatterning of Quantum Dots for Biofunctionalization and Nanoimaging. ACS Appl Nano Mater (2023) doi:10.1021/acsanm.3c00778.

56. Pepelanova, I., Kruppa, K., Scheper, T. & Lavrentieva, A. Gelatin-methacryloyl (GelMA) hydrogels with defined degree of functionalization as a versatile toolkit for 3D cell culture and extrusion bioprinting. Bioengineering 5, (2018).

57. Pamplona, R. et al. Tuning of Mechanical Properties in Photopolymerizable Gelatin-Based Hydrogels for In Vitro Cell Culture Systems. ACS Appl Polym Mater 5, 1487–1498 (2023).

58. Bupphathong, S. et al. Gelatin Methacrylate Hydrogel for Tissue Engineering Applications—A Review on Material Modifications. Pharmaceuticals 15, 171 (2022).

59. Yin, J., Yan, M., Wang, Y., Fu, J. & Suo, H. 3D Bioprinting of Low-Concentration Cell-Laden Gelatin Methacrylate (GelMA) Bioinks with a Two-Step Cross-linking Strategy. ACS Appl Mater Interfaces 10, 6849–6857 (2018).

60. Young, A. T., White, O. C. & Daniele, M. A. Rheological Properties of Coordinated Physical Gelation and Chemical Crosslinking in Gelatin Methacryloyl (GelMA) Hydrogels. Macromol Biosci 20, (2020).

61. Levalley, P. J. et al. Fabrication of Functional Biomaterial Microstructures by in Situ Photopolymerization and Photodegradation. ACS Biomater Sci Eng 4, 3078–3087 (2018).

62. Zhu, M. et al. Gelatin methacryloyl and its hydrogels with an exceptional degree of controllability and batch-to-batch consistency. Sci Rep 9, 1–13 (2019).

63. Guimarães, C. F., Gasperini, L., Marques, A. P. & Reis, R. L. The stiffness of living tissues and its implications for tissue engineering. Nat Rev Mater 5, 351–370 (2020).

64. Raghavan, S., Nelson, C. M., Baranski, J. D., Lim, E. & Chen, C. S. Geometrically controlled endothelial tubulogenesis in micropatterned gels. Tissue Eng Part A 16, 2255–2263 (2010).

65. Jiang, L. Y. & Luo, Y. Guided assembly of endothelial cells on hydrogel matrices patterned with microgrooves: A basic model for microvessel engineering. Soft Matter 9, 1113–1121 (2013).

66. Leslie-Barbick, J. E., Shen, C., Chen, C. & West, J. L. Micron-scale spatially patterned, covalently immobilized vascular endothelial growth factor on hydrogels accelerates endothelial tubulogenesis and increases cellular angiogenic responses. Tissue Eng Part A 17, 221–229 (2011).

67. Haase, K. & Kamm, R. D. Advances in On-Chip Vascularization. Regenerative Med 12, 285–302 (2017).

68. Lehmann, M. et al. Platelets Drive Thrombus Propagation in a Hematocrit and Glycoprotein VI– Dependent Manner in an In Vitro Venous Thrombosis Model. Arterioscler Thromb Vasc Biol 38, 1052–1062 (2018).

69. Sanchez, Z. A. C., Vijayananda, V., Virassammy, D. M., Rosenfeld, L. & Ramasubramanian, A. K. The interaction of vortical flows with red cells in venous valve mimics. Biomicrofluidics 16, (2022).

70. Baksamawi, H. A., Alexiadis, A., Vigolo, D. & Brill, A. Platelet accumulation in an endothelium-coated elastic vein valve model of deep vein thrombosis is mediated by GPIbα—VWF interaction. Front Cardiovasc Med 10, (2023).

71. Baksamawi, H. A., Ariane, M., Brill, A., Vigolo, D. & Alexiadis, A. Modelling particle agglomeration on through elastic valves under flow. ChemEngineering 5, (2021).

72. Schofield, Z., et al. The role of valve stiffness in the insurgence of deep vein thrombosis. Commun Mater 1, (2020).

73. Rajeeva Pandian, N. K., Walther, B. K., Suresh, R., Cooke, J. P. & Jain, A. Microengineered Human Vein-Chip Recreates Venous Valve Architecture and Its Contribution to Thrombosis. Small 16, (2020).

74. Caggiati, A., Phillips, M., Lametschwandtner, A. & Allegra, C. Valves in Small Veins and Venules. European Journal of Vascular and Endovascular Surgery 32, 447–452 (2006).

75. Phillips, M. N., Jones, G. T., Van Rij, A. M. & Zhang, M. Micro-Venous Valves in the Superficial Veins of the Human Lower Limb. Clinical Anatomy 17, 55–60 (2004).

76. Orlova, V. V. et al. Generation, expansion and functional analysis of endothelial cells and pericytes derived from human pluripotent stem cells. Nat Protoc 9, 1514–1531 (2014).

77. Orlova, V. V. et al. Functionality of endothelial cells and pericytes from human pluripotent stem cells demonstrated in cultured vascular plexus and zebrafish xenografts. Arterioscler Thromb Vasc Biol 34, 177–186 (2014).

78. Nahon, D. M. et al. Self-assembling 3D vessel-on-chip model with hiPSC-derived astrocytes. Stem Cell Reports (2024) doi:10.1016/j.stemcr.2024.05.006.

79. Tinevez, J.-Y. et al. TrackMate: An open and extensible platform for single-particle tracking. Methods 115, 80–90 (2017).

80. Ershov, D. et al. TrackMate 7: integrating state-of-the-art segmentation algorithms into tracking pipelines. Nat Methods 19, 829–832 (2022).

